# TREX1 restricts CRISPR-Cas9 genome editing in human cells

**DOI:** 10.1101/2022.12.12.520063

**Authors:** Mehmet E. Karasu, Eléonore Toufektchan, John Maciejowski, Jacob E. Corn

## Abstract

CRISPR-Cas mediated homology-directed repair (HDR) can flexibly introduce desired mutations at targeted sites in a genome. But achieving high HDR efficiencies is a major hurdle in many cellular contexts. Moreover, cells from patients with mutations in DNA repair factors can exhibit low CRISPR-Cas-mediated HDR, complicating genome editing as a potential treatment. We used genome-wide screening in Fanconi anemia (FA) patient lymphoblastic cell lines to uncover suppressors of CRISPR-Cas mediated HDR. Surprisingly, we found that a single exonuclease called *TREX1* is an important determinant of HDR efficiency when single-stranded templates are used as a repair template. *TREX1* expression acts as a biomarker for CRISPR-Cas mediated HDR, such that cell lines expressing high levels of *TREX1* have poor HDR that can be rescued by *TREX1* removal. CRISPR-Cas mediated HDR can also be rescued by using single-stranded DNA templates that are chemically protected in a manner consistent with TREX1’s exonucleolytic activity. Overall, our data provide a mechanistic explanation for why some cells are easier to edit than others and suggest a route to increase CRISPR-Cas mediated HDR in *TREX*-expressing context.

## Introduction

CRISPR-Cas genome editing is a revolutionary technology to introduce targeted mutations in cells and organisms. In its most simple form, Cas enzymes (such as Cas9) are guided by a single guide RNA sequence (sgRNA) to introduce a double stranded break (DSB) at a target genomic locus^1^. This DSB is repaired by one of two main mechanisms: non-homologous end joining (NHEJ) or homology-directed repair (HDR). NHEJ leads to short insertions and deletions (indels) and can be used to disrupt sequences. However, CRISPR-Cas mediated HDR can copy from an exogenously supplied DNA template to introduce sequence changes ranging from single nucleotide polymorphisms (SNPs) to large insertions (e.g. chimeric antigen receptors)^2,3^. The incredible flexibility of HDR makes it attractive for both research and therapeutic applications of genome editing. But in most human cells the efficiency of HDR is very low relative to NHEJ^4,5^.

Cellular context plays a significant role in the efficiency of HDR. Fundamental studies of DNA repair have revealed that HDR is cell cycle limited, and most active in S/G2 ^6–8^. Differential expression, cellular background, and/or mutational burden can also dramatically affect efficiency. CRISPR-Cas9 mediated editing the same locus with the identical reagents in various cell types can result in variable efficiency, ranging from 30% of alleles to complete inactivity^9^. Recent work has shown that prime editing efficiency is determined by mismatch repair status of the targeted cells, yet a similar biomarker for CRISPR-Cas9 mediated HDR efficiency is currently unknown^10,11^.

Patient cells carrying mutations in DNA repair genes can also compromise CRISPR-Cas mediated genome editing, complicating efforts to correct the targeted disorder. For example, loss of function in one of 22 genes involved in Fanconi anemia (FA) can largely prevent HDR ^9,12,13^. FA is a rare genetic disorder characterized by bone marrow failure and predisposition to malignancies later in life^14^. Attempts to correct FA patient mutations by CRISPR-Cas9 induced HDR have revealed poor efficiencies, limiting potentially curative genome editing approaches to those that circumvent HDR but are not applicable to all FA alleles^15,16^.

Here, we performed genome-wide screening in lymphoblastic cells derived from FA patients to uncover factors that restrict CRISPR-Cas9 mediated HDR. We found that TREX1, a widely expressed endoplasmic reticulum (ER)-associated nuclease involved in innate immunity, plays a dominant role in reducing CRISPR-Cas9 induced HDR in human cells. Knockout of *TREX1* rescues HDR in FA patient-derived cells and in commonly-used cell models with naturally low HDR efficiency.Chemical protection of DNA donor templates in a manner designed to prevent TREX1 activity rescues HDR in these TREX1-expressing cells at multiple loci. Our work highlights the importance of cellular factors in regulating genome editing outcomes, provides a rational explanation to the seemingly stochastic efficiency of CRISPR-Cas9 mediated HDR in various cell backgrounds, and offers a potential path to high levels of genome editing in the myriad cell models limited by TREX1 expression.

## Results

### Removal of TREX1 reactivates homology directed repair in Fanconi anemia patient cells

We first analyzed CRISPR-Cas9 mediated HDR efficiencies of LCLs from *FANCA*^*–/–*^ and *FANCD2*^*–/–*^ patient backgrounds with a previously published BFP to GFP reporter ^9,17,18^. This BFP sequence can be targeted by a Cas9 RNP and appropriate single-stranded oligodeoxynucleotide HDR template (hereafter ssODN) to convert BFP-His151 to GFP-Tyr151.^18^ After optimizing electroporation conditions in healthy donor LCLs (**Supplementary figure 1**), we verified that HDR efficiencies in FA LCLs were extremely low as compared to wild-type LCLs, especially in the *FANCA*^*–/–*^ background (e.g. 0.346 ± 0.236% *FANCA*^*–/–*^ vs 2.308 ± 0.586% *FANCD2*^*–/–*^ vs 9.13 ± 0.714% wild type) (**Figure 1A**). We therefore searched for factors whose removal could rescue HDR in *FANCA*^*–/–*^ cells.

**Figure 1:**
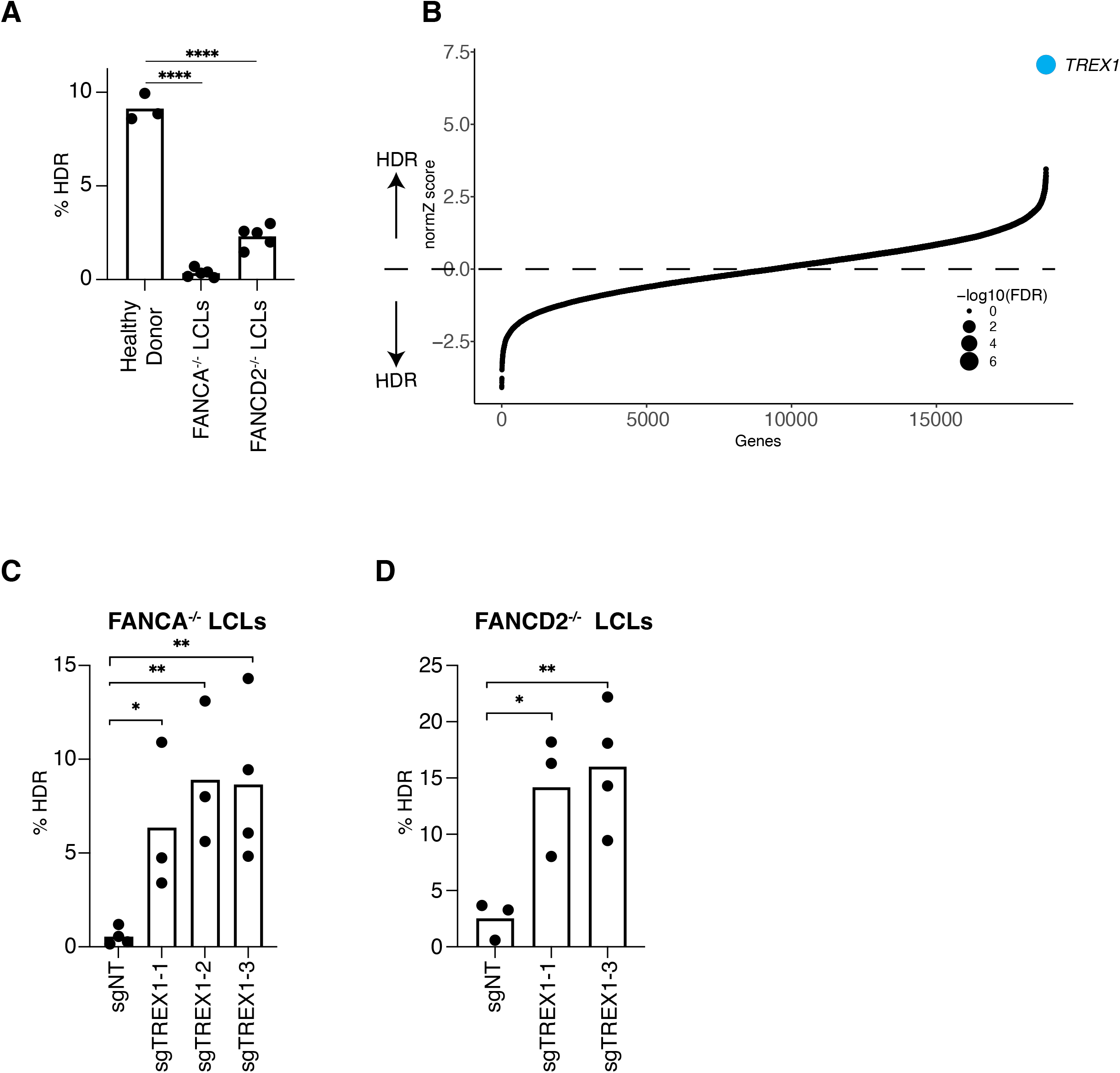
Identification of TREX1 as a restrictive factor of CRISPR-Cas9 mediated HDR. (A) FA patient derived LCLs are compromised in a BFP to GFP reporter assay for CRISPR-Cas mediated HDR. The efficiency of HDR was measured by flow cytometry 5 days after Cas9 targeting. (B) Genome-wide CRISPRi/CRISPRn screening identifies *TREX1* as the sole gene whose knockdown strongly rescues HDR in *FANCA*-/-LCLs. Gene-level effects and statistics were calculated using DrugZ and ranked by the normZ score. The size of each point reflects the false discovery rate. C,D) CRISPRi knockdown of *TREX1* significantly increases CRISPR-Cas9 mediated HDR in *FANCA* (C) and *FANCD2* (D) deficient LCLs. Both cell backgrounds were stably transduced with up to three different sgRNAs targeting *TREX1* and a BFP-to-GFP assay was used to measure HDR efficiency. In A and D) Each dot represents the individual biological replicate measurements and and bars represent the mean. All p-values were calculated using an un-paired t test, * p<0.05, **p<0.01, ***p<0.001.

We used pooled genetic screening to uncover factors regulating CRISPR-Cas9 mediated HDR^17,19,20^. We employed a previously published BFP-to-GFP HDR reporter and paired CRISPR inhibition (CRISPRi) and CRISPR ribonucleoprotein (RNP) system to knock down one gene per cell, while simultaneously inducing a DSB and paired ssODN repair template at the reporter locus ^9,21^. Fluorescence activated cell sorting (FACS) for conversion of BFP to GFP and sequencing of the recovered sgRNAs is used to reveal factors that modulate HDR. Technical concerns limited our prior work in using this screening system to a set of 2,000 core genes involved in DNA metabolism^9^. Here, we developed approaches that allowed scaling up 10-fold to genome-wide screening (see Methods).

*FANCA*^*–/–*^ LCLs were first lentivirally engineered to stably express KRAB-dCAS9-mCherry (CRISPRi). We recloned all guide RNAs in the CRISPRiv2 library ^22^ to a modified reporter lentiviral vector with BFP placed downstream of the sgRNA cassette. During screening, electroporation of Cas9 RNP and ssODN into *FANCA*^*–/–*^ CRISPRi LCLs yielded a GFP-positive fraction of approximately 0.5% (**Supplementary figure 2**). We used a dual thresholding- and pure-sort strategy to rapidly yet accurately isolate the rare cells that were performing high levels of HDR (see Methods). We amplified sgRNAs by PCR from the unsorted and GFP-positive populations after biological duplicate viral transductions, sequenced the population of sgRNAs using next generation sequencing, quantified guide abundances using MaGeck ^23^, and calculated gene-level enrichment scores and significance using DrugZ ^24^. Surprisingly, we found that a single gene called *TREX1* was highly enriched in the GFP-positive HDR population (FDR < 1×10^−7^)) (**Figure 1B, Supplementary Table 1)**.

### TREX1 destabilizes unprotected HDR templates

TREX1 is a 3′-to-5′ exonuclease that is anchored to the outer membrane of the ER and is involved in suppressing chronic activation of cyclic GMP-AMP synthase (cGAS) during the innate immune response to cytosolic DNA ^25–27^. TREX1 is active on single-stranded and double-stranded DNA molecules and has a 1000-fold lower activity toward RNA and RNA-DNA hybrids ^25,28^. Mutations associated with TREX1 lead to auto-immune disorders such as Aicardi-Goutieres syndrome^29–31^. Notably, neither TREX2 (46% identical to TREX1 in the catalytic domain) nor the hundreds of other nucleases present in human cells were screening hits.

To validate the primary screen result, we individually cloned multiple guide RNAs against *TREX1* and performed individual CRISPRi in both *FANCA*^*–/–*^ and *FANCD2*^*–/–*^ LCLs. Quantification of baseline expression of *TREX1* by qRT-PCR revealed very high expression in both FA backgrounds relative to healthy donor cells (**Supplementary figure 3**). CRISPRi knockdown was effective with multiple *TREX1*-targeting guide RNAs. Individual BFP-to-GFP reporter assays showed that *TREX1*-knockdown restored HDR activity in both *FANCA*^*–/–*^ and *FANCD2*^*–/–*^ patient backgrounds (**Figure 1C, Supplementary figure 4**).

Given TREX1’s known activity as an exonuclease, we hypothesized that it might be degrading the DNA repair template used during CRISPR-Cas9 mediated HDR. To test this, we used CRISPR-Cas to create an isogenic *TREX1* knockout in RPE1 cells, which have otherwise functional DNA repair ^32^. We then ectopically expressed wild type TREX1 in the knockout clone ^32^ (**Supplementary Figure 5**). A BFP to GFP assay showed that wildtype RPE1 cells perform moderate levels of CRISPR-Cas9 induced HDR, *TREX1* knockout strongly increased increased HDR, and overexpression of wild type *TREX1* in the knockout background almost completely abrogated HDR. (**Figure 2A**).

**Figure 2:**
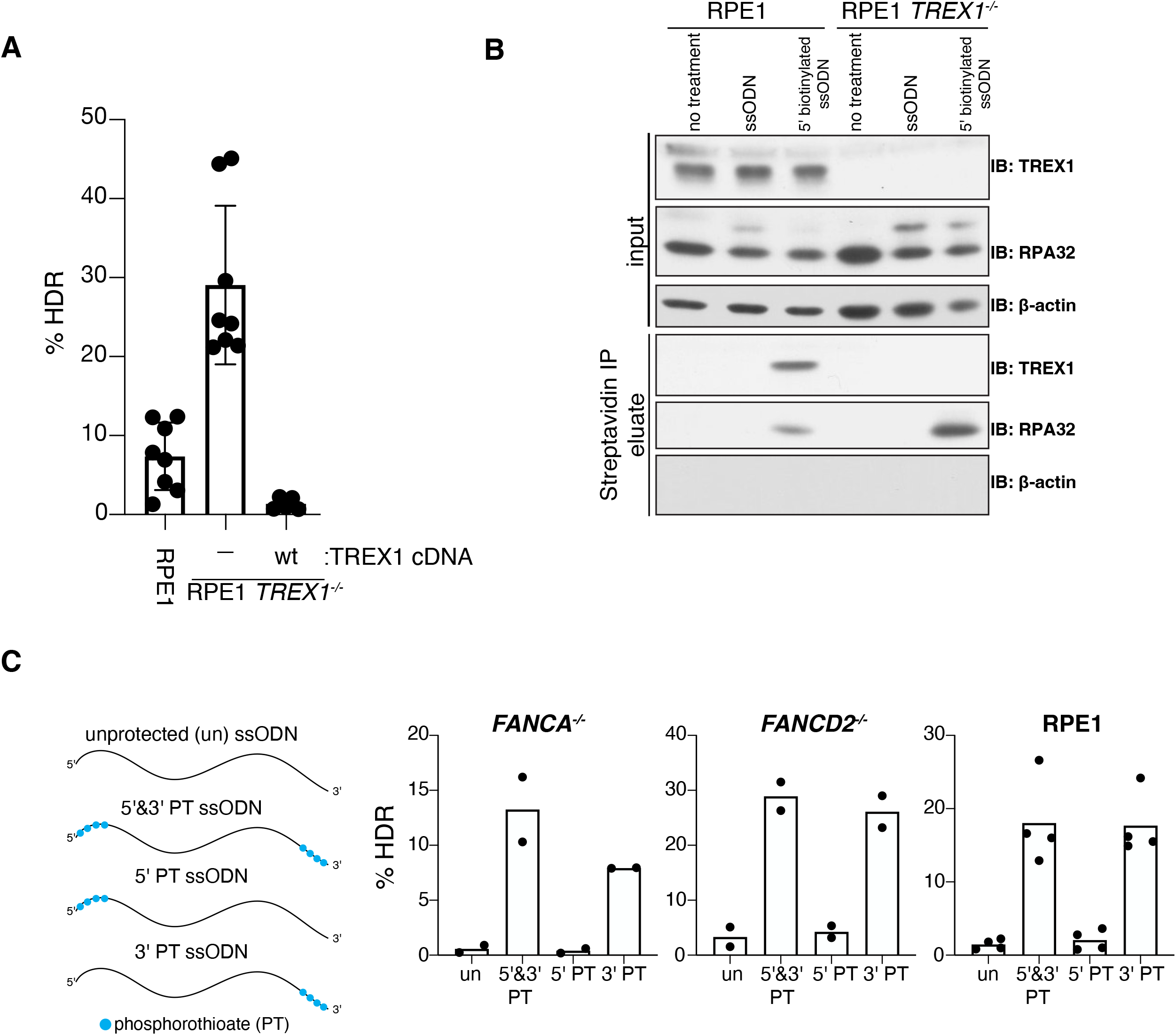
TREX1 interacts with ssODN HDR templates and is inhibited by phosphorothioate protection. (A) RPE1 *TREX1*^*–/–*^cells perform high levels of HDR and *TREX1* cDNA complementation abrogates HDR as measured by the BFP-to-GFP assay. Dots represent individual biological replicate measurements, bars represent the mean values, and error bars represent the standard deviation. (B) TREX1 co-immunoprecipitates with 5’-biotin labeled ssODN template. HDR donors were delivered by electroporation to RPE1 wild type and RPE1 *TREX1*^*–/–*^ cells. After 20 minutes, cells were collected and lysates were prepared for immunoprecipitation with streptavidin beads. Blots were probed with anti-TREX1, anti-RPA32 and anti-β actin. (C) Incorporation of four phosphorothioate (PT) bonds on the 5′ & ′ or only 3’ ends of an ssODN rescues HDR efficiency in *FANCA*^*–/–*^ LCLs, *FANCD2*^*–/–*^ LCLs, and RPE1 cells. 5’ end protection behaves as unprotected (un) ssODN. HDR was measured using the BFP-to-GFP assay. Dots represent the individual biological replicate measurements and bars represent the mean.

Next, we asked whether TREX1 physically interacts with an ssODN delivered to cells. We electroporated *TREX1* wild type and *TREX1*^*–/–*^ RPE1 cells with a 5’ biotinylated ssODN template and performed immunoprecipitation using streptavidin beads 2 hours after nucleofection. As a positive control, we blotted for RPA32, which interacts strongly with ssODNs ^33^. RPA32 was readily measurable in the immunoprecipitated samples from both *TREX1* wild-type and *TREX1*^*–/–*^ cells. *TREX1* was also strongly associated with the biotinylated ssODN in the *TREX1* wild type sample, demonstrating that TREX1 forms a stable interaction with electroporated ssDNA templates (**Figure 2B**).

If TREX1 is indeed a prominent exonuclease affecting the stability of ssODN templates, a 3’ protected ssODN should block its 3’-to-5’ hydrolytic activity ^34^. We therefore measured HDR efficiency using protected ssODNs that have phosphorothioate bonds between five nucleotides at the 5’- and/or 3’-ends (**Figure 2C**). An unprotected ssODN yielded moderate HDR in wild type RPE1 cells and very low HDR in *FANCA*^*–/–*^ and *FANCD2*^*–/–*^ LCLs. Protection of the ssODN at both the 5’ and 3’ ends markedly increased HDR in all cell types. Consistent with the polarity of TREX1 activity, individual 3’ end protection was sufficient to increase HDR, whereas 5’ end protection performed similar to the unmodified ssODN.

### TREX1 expression predicts HDR efficacy in multiple cell lines and ssODN protection rescues HDR in TREX1-expressing contexts

We wondered if TREX1 could explain the widely variable CRISPR-Cas9 mediated HDR efficacy observed in different human cell types. *TREX1* expression is reported to differ widely between cell lines, which we confirmed via qRT-PCR (**Supplementary Figure 6-7**). Cell lines commonly used for genome editing with anecdotally high levels of HDR, including K562 and HEK293, have low levels of *TREX1*. By contrast, cells where HDR is anecdotally much more difficult, such as U2OS and Jurkat, express high levels of *TREX1*.

To determine whether *TREX1* expression is a predictive marker for CRISPR-Cas mediated HDR efficiency, we tested an integrated BFP-to-GFP reporter in Jurkat, MDA-MB-321, U2OS, HeLa, and K562 cells. HeLa cells are one of the highest *TREX1*-expressing cell lines according to the Human Protein Atlas (https://www.proteinatlas.org/), and we found that they exhibit extremely low basal CRISPR-Cas9 mediated HDR. Importantly, HeLa HDR efficiency can be rescued to the same level as K562 cells by CRISPRi knockdown of *TREX1* or use of a protected ssODN (**Supplementary Figure 8**). Among the cell backgrounds we tested, only *TREX1*-low K562 cells exhibited high HDR with an unprotected ssODN (**Figure 3A**). Protected ssODNs increased CRISPR-Cas9 HDR in all *TREX1*-expressing cells tested, but did not further increase HDR in *TREX1*-low K562s (**Figure 3A**).

**Figure 3:**
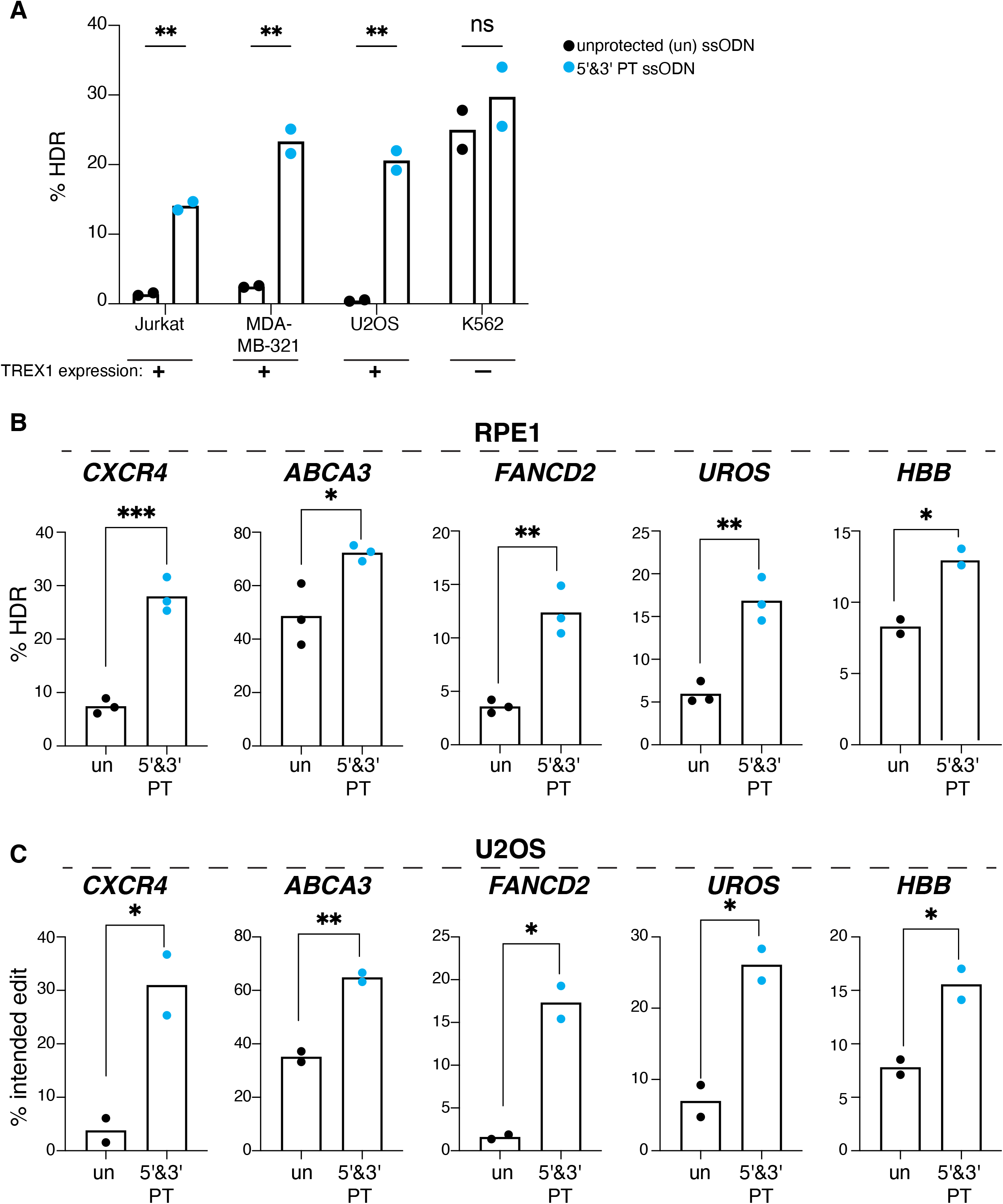
Protected ssODN templates increase HDR in *TREX1*-expressing cell contexts at multiple loci. A) Cell types expressing *TREX1* exhibit compromised HDR, but this is rescued by using phosphorothioate protected ssODNs. A cell type with normally low levels of *TREX1* is already efficient at CRISPR-Cas HDR and this is not further improved by a protected ssODN. HDR was measured by the BFP-to-GFP assay in Jurkat, MDA-MB-321, U2OS and K562 cell lines. Black dots represent measurements with unprotected (un) ssODN template and blue dots represent measurements with 5′ & ′ protected (PT) ssODN templates. Each dots represent individual biological replicate measurements. B,C) CRISPR-Cas mediated HDR efficiency is rescued at multiple endogenous loci in *TREX1*-expressing RPE1 cells (C) and U2OS cells (D) by phosphorothioate ssODN protection. Editing sites and HDR mutations are shown in **Supplementary Figure 9**. Black dots indicate use of unprotected (un) ssODN templates and blue dots represent 5′ & ′ PT ssODN templates. All p-values were calculated using un-paired t test, * p<0.05, **p<0.01, ***p<0.001.

We finally tested the effect of ssODN protection at introducing multiple mutation types (single nucleotide changes, short insertions, and short deletions) at multiple endogenous loci in two different *TREX1*-expressing cells with normally low CRISPR-Cas mediated HDR (RPE1 and U2OS cells) (**Supplementary figure 9**)^3,35,36^. RPE1 and U2OS cells were edited using Cas9 RNPs and either protected or unprotected ssODNs, with editing efficiencies measured by NGS. Using unprotected ssODNs, we found relatively low HDR at almost every locus in both cell types. Phosphorothioate protection increased HDR efficiency by 1.8 to 3.7 fold at every locus tested in both RPE1 and U2OS (**Figure 3B-C**). A protected ssODN also rescued HDR efficiency at *UROS* in HeLa cells, a line with among the highest *TREX1* expression in the Human Protein Atlas (**Supplementary Figure 10**). Taken together, our data indicate that *TREX1* is a major restrictor of CRISPR-Cas mediated HDR in human cells, which can be circumvented by either removing *TREX1* or using DNA templates that are chemically protected from TREX1 (**Figure 4**).

**Figure 4:**
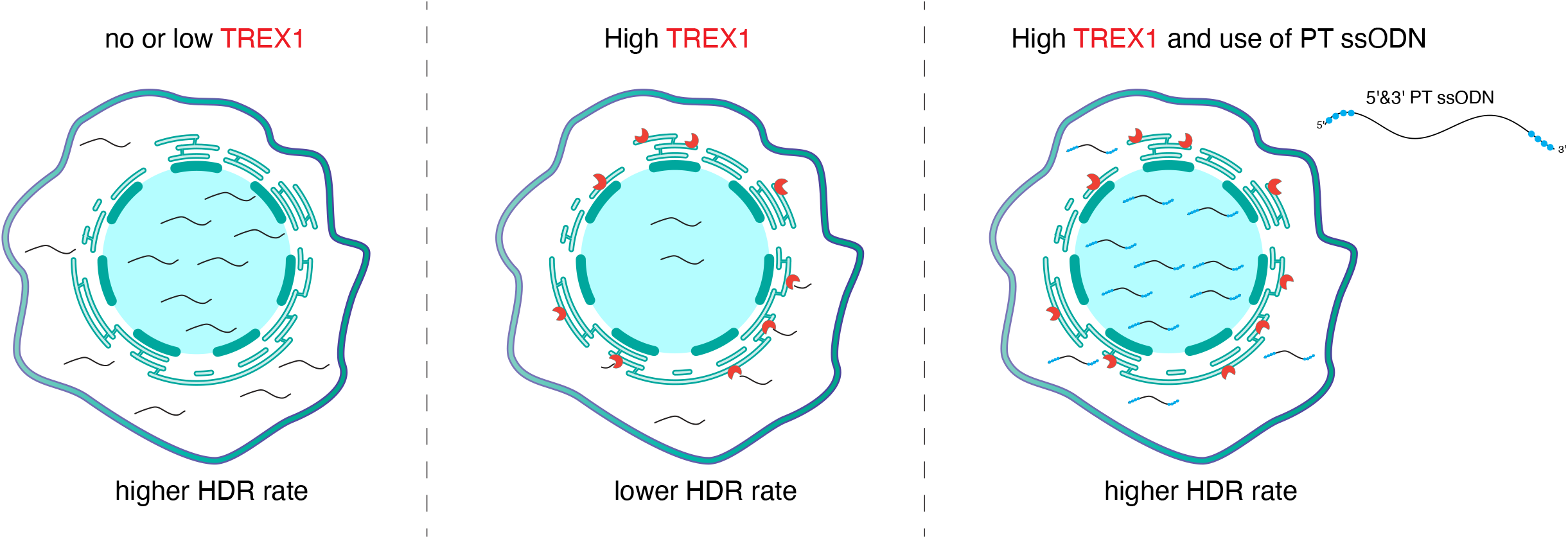
A model for TREX1’s role in suppressing CRISPR-Cas HDR. In *TREX1* low cells, ssODN templates are abundant throughout the cell and available in the nucleus for efficient CRISPR-Cas mediated HDR. In cases where *TREX1* expression is high, ssODN templates are 3’-to-5’ degraded by TREX1 at the endoplasmic reticulum. This reduces ssODN availability in the nucleus and reduces HDR efficiency. Phosphorothioate protection prevents TREX1 digestion of the ssODN, maintaining a high template concentration throughout the cell and enabling efficient CRISPR-Cas mediated HDR.

## Discussion

We identified TREX1 as an exonuclease that restricts CRISPR-Cas9 mediated HDR in human cell lines. Cell lines expressing low levels of TREX1 are competent to perform HDR and have become the cell backgrounds where a great deal of work using CRISPR-Cas HDR has been done. By contrast, FA cells and multiple immortalized cell lines express high levels of TREX1 and are compromised in their ability to perform HDR. Removal of TREX1 from these cells markedly increases CRISPR-Cas HDR, as does the use of chemically protected ssODN templates that would specifically escape TREX1 activity. The presence of TREX1 as the sole hit during screening and the ability of TREX1 removal alone to rescue HDR to high levels suggests that, in cells normally transiting through the cell cycle, it may be a dominant factor involved in restricting CRISPR-Cas HDR from exogenously provided templates.

As an ER-resident enzyme ^25,32,37^, TREX1 probably degrades cytoplasmic DNA templates prior to their diffusion into the nucleus. However, some studies have suggested that TREX1 may be actively involved in DNA repair and could shuttle to the nucleus after DNA damage ^25,38^. Our data using protected ssODNs indicates that TREX1’s main role in restricting CRISPR-Cas mediated HDR is related to template availability. But it is possible that TREX1 transits to the nucleus after Cas-induced damage to also degrade nuclear templates. Further microscopy studies could address exactly where TREX1 degrades the majority of DNA templates during genome editing.

Phenomenological studies have previously hinted at the benefits of shielding the ends of HDR templates during genome editing. Dual 5’ and 3’ phosphorothioate protection of an ssODN can increase HDR in some contexts ^39,40^. But a lack of mechanistic understanding meant that the ideal situations to protect with phosphorothioates were unknown, and they have not been generally adopted by the field. ssODNs can also be protected by integrating Cas9 binding sites at their termini or by circularizing the single stranded template ^41,42^. We propose that all of these protections at least partly work by protecting against TREX1 activity, and their benefits would be even further enhanced in cell backgrounds with high levels of TREX1. We suggest *TREX1* expression as a biomarker for the use of ssODN protection, and that 3’ end protection is sufficient to protect against TREX1 activity. Small molecule inhibition of TREX1 during genome editing might also increase HDR and could be useful if phosphorothioate protected templates are toxic to a certain cell type ^43^. But TREX1 inhibitors are not broadly available and would be more complicated to use than protection of the DNA template.

Overall, our data shed mechanistic light on why donor template protection increases HDR, provide a concrete biomarker for the targeted use of template protection, and resolve long-standing confusion around why some cell types are easier to edit than others.

## Methods

### Cell lines

Healthy donor, *FANCA*^*–/–*^ (FA55) and *FANCD2*^*–/–*^ (FA-75) lymphoblastic cell lines (LCLs) were generously gifted by Dr. Paolo Rio, CIEMAT. K562, Jurkat, U2OS, MD-MBA-431, RPE1, HELA were obtained from ATCC or Berkeley Cell Culture facility. RPE1 *TREX1*^*-/-*^ cell line was used in this paper ^32^. LCLs were cultured in Roswell Park Memorial Institute medium (RPMI 1640, GlutaMAX™, from Thermo Fisher Scientific, 61870010) supplemented with 20% Gibco™ fetal bovine serum (FBS) (ThermoFisher Scientific, 10270106), 1% Gibco™ penicillin/streptomycin (P/S) solution (ThermoFisher Scientific, 15140122), 0.005CmM Gibco™ β-mercaptoethanol (ThermoFisher Scientific, 31350010) and 1% Gibco™ MEM non-essential amino acids (ThermoFisher Scientific, 11140050). K562 and Jurkat cells were cultured in RPMI 1640 GlutaMAX™ media supplemented with 10% FBS and 1% P/S solution. U2OS, MD-MBA-421, RPE1, HELA were cultured in DMEM, high glucose, GlutaMAX™ pyruvate medium (ThermoFisher Scientific, 10569010) supplemented with 10% FBS and 1% P/S. All cells were cultured at 37 °C with 5% CO_2_ in a humidified incubator. Cell lines were regularly tested for mycoplasma with MycoAlert Mycoplasma Detection Kit (Lonza, LT07-318).

To generate CRISPRi cell lines, we packaged a CRISPR inhibition (pHR-EF1a-dCas9-HA-mCherry-KRAB-NLS) construct into lentiviruses in HEK293T cells. The lentiviral supernatant was filtered and used to transduce LCLs and HELA cells. After transduction, mCherry positive cells were sorted using SH800 Cell sorter (Sony). Since LCLs failed to survive as single cells in 96-well plates, we first seeded ∼500 mCherry negative LCLs per well in 96-well plates, then sorted single mCherry positive cells into 500 mCherry negative cells to overcome the viability problem. mCherry positive LCLs were further enriched with consecutive rounds of sorting until reaching mCherry purity >90%.

For retroviral transduction of 3×flag-TREX1-wt in RPE-1 hTERT TREX1-KO cells, open reading frames were cloned into pQCXIZ, which confers resistance to zeocin. Constructs were transfected into Phoenix amphotropic packaging cells using calcium phosphate precipitation. Cell supernatants containing retrovirus were filtered, mixed 1:1 with target cell media and supplemented with 4 μg /ml polybrene. Successfully transduced cells were selected using zeocin (Life Technologies).

### In vitro transcription of gRNAs

gRNAs were in vitro transcribed as described (dx.doi.org/10.17504/protocols.io.dwr7d5) ^44^. Briefly, overlapping oligomers, indicated in the **Supplementary Table 2**, containing a T7 promoter, protospacer and gRNA scaffold were amplified by Q5 High-Fidelity polymerase (New England Biolabs, M0491L) for 15 cycles. 1 µM of T7FwdLong and T7RevLong were used as a template and amplified by T7FwdAmp and T7RevAmp in 50 µl reaction volume. 8 µl PCR amplified product was used for the in vitro transcription using the NEB HiScribe T7 High Yield RNA Synthesis kit (New England Biolabs, E2040S), incubating at 37 °C for 18 hours in the thermocycler. Then, the reaction was supplemented with DNase I (Qiagen, 79256) for 30 mins at 37 °C, followed by Quick CIP (New England Biolabs, M0525S), treatment for 1 hour at 37 °C. The gRNAs were later purified with miRNeasy kit (Qiagen, 217604), concentration was measured by Qubit™ RNA Broad Range (BR) assay (ThermoFisher Scientific, Q10211) and stored at -80 °C.

### Genome-wide library construction

To shuttle genome-wide sgRNAs, we first amplified the cassettes including sgRNA using the primer set priEK-35 and priEK-37 from the CRISPRi-V2 library (Addgene #1000000093) using Phusion polymerase (New England Biolab, M0530L) under the following condition: 30 sec at 98 °C, then 15 cycles of 15 sec at 98 °C, 15 sec at 53 °C, 15 sec at 72 °C, then a final extension for 10 min at 72 °C. Amplified PCR fragments were digested overnight with Bpu1102I (BlpI) (ThermoFisher Scientific, ER0091) and BstXI (ThermoFisher Scientific, ER1021) at 37 °C. The digested DNA fragments were separated 10% TBE gel to cut the DNA band corresponding (∼33 bp). The gel pieces were crushed by spinning for 3 mins at 20k × g and then eluted in water at 37 °C overnight. DNA later was precipitated with NaOAc/ EtOH method. Meanwhile, we linearized the vector carrying mutated GFP sequence the same restriction enzymes, Bpu1102I (BlpI) and BstXI for 4 hours at 37 °C. The linearized DNA product was separated by 0.8% agarose gel electrophoresis and excised from the gel. DNA was cleaned by QIAquick Gel extraction kit (Qiagen). The DNA further was cleaned with NaOAc/ EtOH method. For the ligation reaction, 500 ng linearized vector and 1.9 ng insert was incubated with T4 DNA Ligase (New England Biolab, M0202L) for 16 hours at 16 °C. The ligated plasmids were purified by isopropanol/ 5M NaCl precipitation and resuspend in 13 µl elution buffer. 1 or 2 µl of the purified ligation reaction were mixed with 25 µl of MegaX DH10B™ T1^R^ electrocompetent cells (ThermoFisher Scientific, C640003) and recovered in S.O.C medium for 1.5 hours. The bacteria were plated in 24.5 × 24.5 cm LB Agar plates containing ampicillin resistance. While plating the bacteria in 24.5 × 24.5 cm, the dilutions of bacteria were performed as well to detect approximately coverage of sgRNA library. Grown colonies were collected by scraping from LB-Agar plates. Plasmids were recovered by several midi-preps (Qiagen Plasmid Plus Midi Kit).

Quality of the library was determined by next generation sequencing. Sequencing libraries were prepared by amplifying sgRNA cassettes with the primers priEK_i5-1 and priEK_i7-1, and secondary PCR to put the sequencing adapters priEK_501-priEK_701 (indicated in the supplementary file). The reaction was sequenced by MiSeq and sgRNA distribution of the cloned library was analyzed using custom scripts.

### Production of sgRNA library lentiviruses

To produce lentivirus from the sgRNA library, ∼ 7 million HEK293T cells were seeded in a 15 cm plate in 20 ml of DMEM medium with 10% FBS and 1% P/S. The following day HEK293T cells were transfected with the library. Per plate, in a 5 ml tube, 15 µg of sgRNA library, 12 µg of delta VPR and 3 µg of VSVG were resuspended with 1.3 ml OPTIMEM and mixed with 270 µl polycation polyethylenimine 1 mg/ml (PEI, 1 µl to 3 µg of DNA). The mix was incubated at room temperature for 20 min and then added on top of HEK293T cells in a drop-wise manner. The media was changed on the following day. The virus-containing cell culture media was collected 48 hours and 72 hours after the transfection. The viral media was combined and filtered using 0.45 µm PES membrane (ThermoFisher Scientific, 295-3345), aliquoted into 15 ml Eppendorf tubes, snap frozen and stored at -80 °C.

### CRISPR screen

*FANCA*^*–/–*^ CRISPRi cells were grown to 150 million cells before transduction with the genome-wide CRISPRi library. Since LCLs were extremely difficult to transduce with lentiviruses, *FANCA*^*–/–*^ CRISPRi were directly resuspended in virus-containing media and seeded in 6 well plates in the presence of 8 µg/ml polybrene. The coverage determined by BFP positive cells was around 300 × per sgRNA. 24 hours after transduction, cells were collected and transferred to T75 flasks. A day later, guide RNA containing cells were selected with puromycin treatment (0.5 µg/ml) for 96 hours. At this moment, BFP positive cells were over 90%. Upon puromycin selection achieved, cells were split into two replicates and were maintained for 250× coverage throughout the screen. 20 days after transduction, cells were subjected to Ficoll gradient (Ficoll® Paque Plus, Millipore Sigma, GE17-1440-02). 1 × 10^6^ cells were electroporated with 400 pmol SpCas9-nuclear localization sequence (NLS), 480 pmol L2 gRNA targeting BFP and 500 pmol BFP to GFP ssODN template using CM-189 and SF solution (Lonza, 4D electroporator) per replicate. Cells were further expanded in culture prior to sorting. Before the sort, a background population was collected for downstream NGS analysis. The sort was performed in two steps: first thresholding was set to enrich the GFP positive population from (∼ 0.5 % to 70 %) and then a stringent sort was performed to achieve ∼99% pure GFP positive cells. Cells were pelleted and stored at -80 °C until genomic DNA extraction.

### NGS sample preparation and screen analysis

Genomic DNA was extracted using Gentra PureGene Cell Kit (Qiagen, 158912) gDNA extraction protocol. Briefly, for the background samples (from total 25 million cells), cell pellet resuspended in 3 ml cell lysis solution and then mixed with 15 µl RNase A solution at 37 °C for 20 mins, cooled it down for 10 mins on ice and add 1 ml protein precipitation buffer. The mix was vortexed thoroughly and was spun down for 10 mins, 2000 × g. The genomic DNA containing supernatant was mixed with 100% isopropanol (3 ml) by inverting the 15 ml tube for 50 times. The genomic DNA was pelleted by centrifugation at 2000 × g, for 5 mins, and then washed with 70% EtOH. After removing 70% EtOH, genomic DNA was resuspended in 200 µl hybridization buffer, and incubated at 50 °C for 1 hour. DNA amount was measured by Nanodrop. For the GFP positive cells, the protocol was adjusted for the low cell number. In the genomic DNA precipitation step, glycogen was added to facilitate the genomic DNA precipitation.

Purified genomic DNA were used for the further PCR amplification of sgRNA cassettes following the protocol (weissman.wi.mit.edu/resources/IlluminaSequencingSamplePrep.pdf). For the background samples 5 µg genomic DNA per reaction was used. For the sort background samples, 30 PCR reactions were performed and combined later. Since we had very limited DNA from GFP positive cells, we used 1 µg amount of DNA per reaction and performed two reactions. The PCR products were purified by two rounds of Sera-Mag magnetic beads (Cytiva, 29343957). The concentrations were measured by Qubit™ 1 × dsDNA high sensitivity assay (ThermoFisher Scientific, Q33232) and the samples were pooled according to their anticipated read counts. The samples were sequenced on Next Seq 2000.

Screening data were analyzed using the standard protocols in MaGECK and DrugZ. MaGECK was used to get the guide RNA counts per each sgRNA in the population and DrugZ was used to integrate multiple guides into gene-level phenotypes relative to the background unsorted population (normZ score and FDR values)^23,24^ (**Supplementary Table 1**).

### RNP electroporation for BFP to GFP reporter assay and genomic loci targeting

RNP electroporation was performed as described (dx.doi.org/10.17504/protocols.io.dm649d) ^9^. Briefly, 36 pmol sgRNA and 30 pmol SpCas9-NLS were mixed in Cas9 buffer (20 mM HEPES at pH 7.5, 150 mM KCl, 1CmM MgCl_2_, 10% glycerol and 1 mM tris (2-carboxyethyl) phosphine (TCEP) reducing agent). The mixture was incubated at room temperature for 20 mins. Meanwhile, 1 × 10^5^ to 2 × 10^5^ cells were collected and spun down at 300 × g for 5 mins. The cell pellets were resuspended in 15 µL of nucleofection buffer (Lonza). Then 5 µl of RNP mixture was added to the cell suspension with 0.3 µl of 100 µM (30 pmol) ssODN (BFP to GFP template) template. 5 days after electroporation cells were collected and subjected to flow cytometry with an Attune Flow Cytometer (ThermoFisher Scientific). Downstream analysis was performed using FlowJo Software v10.8.2 (FlowJo, LLC)

For endogenous locus targeting, 100 pmol SpCas9-NLS was mixed with 120 pmol gRNA in Cas9 buffer and the mixture were incubated for 20-30 min at room temperature or 37 °C. 1 × 10^5^ to 2 × 10^5^ cells were collected and resuspended in 15 µL of nucleofection buffer (Lonza). For each reaction, 100 pmol of ssODN was then added before nucleofections. Electroporations were performed in the strip format, with 20 µl volume of cells and RNP mix. The following kit and program for each cell type was selected: K-562 (SF kit/FF-120), RPE1 (P3 kit/ EA-104), U2-OS (SE kit/ CM-130), MDA-MB-421 (SE kit/CM-130), HELA (SE kit/ CM-130), Jurkat (SE kit/ CL-120). After electroporation, prewarmed 80 µl of DMEM or RPMI medium was added into strips. Cells were incubated in the hood for 10 mins and then transferred to the plates and returned to 37 °C.

ssODNs were purchased from Integrated DNA Technologies as Ultramer DNA oligos. To protect ssODNs, they were ordered with four phosphorothioate modifications at the 5′ and/or 3′ ends. Sequence information can be found in the **Supplementary table 2**.

### Genomic DNA extraction

Cell pellets were collected 72-96 hours after electroporation and resuspended in QuickExtract solution (Lucigen, QE09050) and subjected to genomic DNA extraction while incubating for 10 min at 65 °C, 5 min 98 °C and then holding at 4 °C. Following the incubation, 1 µl of genomic DNA was taken for further PCR reactions for next generation sequencing.

### Next Generation Sequencing

Primers containing adaptor binding sites (indicated in the **Supplementary Table 2**) were designed to amplify 150-200 bp around the cut sites. First, genomic DNA was amplified using NEBNext^®^ Ultra™ II Q5^®^ Master Mix 30 cycles, and then cleaned with SPRI beads (SeraMag Select (Cytiva, 29343052) or in house). From the purified reactions, around 10-20 ng DNA were used as input for the second PCR reaction to add i7/i5 indexes for the samples in 9 reaction cycles. Amplicons were then purified again with 0.8× SeraMag beads and samples for the same genomic loci were combined. The amplicon length and purity were analyzed by a TapeStation with D1000 DNA flow cells (Agilent). Pools were combined based on their amount and desired read number (50 – 100k reads /per sample). The combined samples were sequenced in Illumina sequencers (MiSeq or NextSeq2000) in the Genome Engineering and Measurement lab at ETH Zurich.

### NGS Analysis

The sequencing reads were demultiplexed and analyzed with CRISPresso2 in batch mode ^45^ with default parameters other than minimum average read quality of 30 and minimum single bp quality of 10. Reads with a frequency lower than 0.5% were disregarded before further analysis. Results were then normalized to sum up to 100%.

### Immunoprecipitation-western blot

Cells indicated in Figure 2 were harvested by trypsinization and lysed in RIPA buffer (5 mM Tris-HCl pH 7.6, 150 mM NaCl, 1% NP-40, 1% sodium deoxycholate, 0.1% SDS), supplemented with phosphatase inhibitors (10 mM NaF, 20 mM β-glycerophosphate) and protease inhibitor (Thermo Scientific) at ∼10^6^ cells/mL and incubated on ice for 20 min. Lysates were sonicated with a Bioruptor Plus sonication device (Diagenode) for 15 cycles ON/OFF (high, 4 °C). Sonicated lysates were incubated on ice for 20 min, centrifuged at 16,000 × g, 4 °C for 20 min and supernatants were transferred into french tubes before protein quantification using the Pierce BCA protein assay (Thermo Fisher Scientific). Lysate equivalent to 10-50 µg of proteins was mixed to 1× Laemmli buffer (50 mM Tris, 10% glycerol, 2% SDS, 0.01% bromophenol blue, 2.5% β-mercaptoethanol), resolved by SDS-PAGE (Life Technologies) and transferred to nitrocellulose membranes (Amersham).Membranes were blocked in 5% milk in TBS with 0.1% Tween-20 (TBS-T) and incubated with primary antibody overnight at 4 °C, washed 3 times in TBS-T, and incubated for 1 hr at room temperature with horseradish-peroxidase-conjugated secondary antibody. After 3 washes in TBS-T, imaging was performed using enhanced chemiluminescence (Thermo Fisher).

### Antibodies

Primary antibodies: anti-TREX1 (abcam ab185228), -ß-actin (abcam ab8224), - RPA32 (abam ab2175) and -flag (abcam f1804).

Secondary antibodies: Goat anti-Mouse IgG HRP (Thermo Fisher Scientific 31432) and Donkey anti-rabbit IgG HRP (SouthernBiotech 6441-05).

### Biotin-ssODN immunoprecipitation

5-6×10^6^ of RPE-1 hTERT parental or TREX1-KO cells expressing were electroporated with 5 nmol biotinylated or unprotected ssODN (IDT) in 100 µL final volume using Lonza 4D-nucleofector (see methods). 2 hrs after electroporation, cells were harvested by trypsinization, washed with PBS and resuspended in lysis buffer (50 mM Tris pH 7.5, 200 mM NaCl, 0.075% NP-40, protease inhibitors) at 10^7^ cells/mL. Cells were then dounce homogenized by 10 strokes with a tight-fitting pestle. Lysates were incubated on ice for 20 min, centrifuged at 16,000 × g, 4°C for 20 min and input samples were taken for immunoblotting. To reduce non-specific binding of proteins to the beads, lysates were precleared by incubation with Protein G dynabeads (invitrogen) for 30 min at room temperature. To pull down the biotinylated ssODN, the cleared lysate was transferred onto streptavidin dynabeads (10-μL per sample, Invitrogen) and again incubated for 30 min at room temperature. The beads were washed 8 times with lysis buffer and then eluted with 2 × Laemmli buffer (100 mM Tris, 20% glycerol, 4% SDS, 0.02% bromophenol blue, 5% β-mercaptoethanol). Immunoblotting of input and eluted samples (diluted 1:2) was performed as described in Methods.

### Quantitative real-time PCR (qRT-PCR)

RNA extraction was performed using RNeasy Mini Kit (Qiagen) according to the manufacturer’s instructions. 1μg of RNA per sample were used for reverse transcription using iScript™ Reverse Transcription Supermix for RT-qPCR (Biorad, 1708841) according to the manufacturer’s instructions. qRT-PCR reactions were set up using SsoAdvanced Universal SYBR Green Supermix (Biorad, 1725271) and run in triplicates using a QuantStudio 6 system (ThermoFisher Scientific). A complete list of primers used in RT-qPCR can be found in the **Supplementary Table 2**.

## Statistical analysis

Each point represents an individual biological replicate and bars represent the mean of the replicates. All p-values were calculated using un-paired t test, * p<0.05, **p<0.01, ***p<0.001 usign Prism GraphPad version 9.4.1.

## Data Availability

Sequencing files for the pooled screen and endogenous genome editing in RPE1, U2OS and HELA cells will be uploaded to GEO/SRA.

## Code Availability

The code used to analyze CRISPR screen and NGS on genomic sites would be available upon request.

## Author Contributions

M.E.K and J.E.C conceived the study. M.E.K designed and performed experiments in figure 1, figure 2 C and figure 3. E.T designed and performed experiments in figure 2A and B. M.E.K and J.E.C wrote the first draft of the paper with input from E.T and J.M.

## Acknowledgements

We thank the Functional Genomics Center Zurich (FGCZ) and especially Dr. Susanne Kreutzer and Dr. Zacharias Kontarakis for their help with NGS sequencing. We thank Dr. Paula Rio and Prof. Jordi Surrallés laboratory (San Pau Hospital, Barcelona) for kindly providing FA-55 and FA-75 LCLs. We thank Dr. Eric J. Aird for kindly providing primers and ssODN templates for *UROS* and *ABCA3* genomic sites and also great inputs during the study and for the manuscript. We thank Lena Kobel for her help handling the various cell lines. We thank Charles Yeh for providing NGS primers for HBB site. We thank Dr.Ana Gvozdenovic for her thoughtful inputs for the manuscript preparation. We also thank the members of the Corn Lab for helpful discussions and help with the manuscript.

JEC is supported by the NOMIS Foundation and the Lotte and Adolf Hotz-Sprenger Stiftung. MEK is supported by the Fanconi Anemia Research Foundation. This project has received funding from the European Research Council (ERC) under the European Union’s Horizon 2020 research and innovation programme (grant agreement No 855741, DDREAMM).

## Supplementary figure legends

**Supplementary figure 1: Optimization of electroporation codes for healthy donor and FA patient derived LCLs**.

BFP to GFP assay was performed in HD and FA LCLs. Indicated codes were tried to find the optimal targeting activity. Targeted cells were gated to measure GFP percentages 5 days after electroporation.

**Supplementary figure 2: Sorting strategy for genome wide CRISPRi strategy**. Representation of *FANCA*^*-/-*^ LCLs sorting in BD FACSDiva 9.0.1. To increase GFP positive cells (originally 0.4%), threshold was set around 10^3^ for the GFP gate and cells were selected according to this threshold (top panel). Later, GFP positive enriched cells (after thresholding sort around ∼ 79 %) were subjected to stringent sort (middle panel). The stringent sort yielded around ∼ 96 % GFP positive cells (bottom panel).

**Supplementary figure 3: RT-qPCR for *TREX1* mRNA levels after CRISPRi depletion**.

RT-qPCR analysis of RNAs extracted from indicated cell lines. The plotted values represent the log_2_ fold difference normalized to healthy donor sample. Two independent experiments were performed, as represented as dots. Bars indicate means for each cell line.

**Supplementary figure 4: Representative flow images for BFP to GFP conversion in *FANCA***^***-/-***^ **LCLs**.

CRISPRi cells were transduced and selected with sgNT (non-targeting), sgTREX1-1 and sgTREX1-3 guide RNAs. Later, BFP to GFP assay was performed in these cells. 5 days after electroporation, GFP was quantified as shown in the flow graphs.

**Supplementary Figure 5: TREX1 protein detection by the Western blotting**. Whole cell extracts were prepared from RPE1 cell lines. The proteins were detected by anti-TREX1, anti-FLAG and anti-ß actin antibodies.

**Supplementary Figure 6: *TREX1* expression dataset from Protein Atlas**. Expression of *TREX1* was searched on Protein Atlas (https://www.proteinatlas.org/) and normalized transcription expression values (nTPM) from various cell lines were replotted. Cell lines used in this study or commonly used in gene editing experiments were highlighted with red.

**Supplementary Figure 7: RT-qPCR for *TREX1* mRNA levels in various cell lines and after CRISPRi depletion in HELA cells**.

RT-qPCR analysis of RNAs extracted from indicated cell lines. The plotted values represent the log_2_ fold difference normalized to K562 cell line. Two independent experiments were performed, as represented as dots. Bars indicate means for each cell line.

**Supplementary Figure 8: Depletion of *TREX1* or using protected ssODN rescue HDR efficiency in HELA cells**.

CRISPRi cell lines were transduced with sgNT (non-targeting) or sgTREX1-3 guide RNAs as indicated. BFP to GFP assay was performed with or without protected ssODN. 5 days after electroporation, GFP was measured and plotted. Two independent experiments were performed (represented as dots).

**Supplementary Figure 9: Endogenous targets and the intended edits for each target**.

For *HBB, CXCR4, FANCD2, UROS* and *ABCA3*, PAM sites are highlighted with dark blue and dashed lines show Cas9 cut site. In the intended edit site, nucleotide changes were highlighted as bold characters, deletions with –. Briefly, *HBB* intended edit is nucleotide substitutions, *CXCR4* and *UROS* intended edits are small inserts, *FANCD2* intended edit is a nucleotide substitution and deletion, *ABCA3* intended edit is two nucleotide small deletion. For *HBB, CXCR4, FANCD2*, the PAM sequences are mutated to prevent for Cas9 recutting.

**Supplementary Figure 10: *UROS* targeting in HELA cell line**.

*UROS* genomic site was chosen to edit by CRISPR-Cas9 and ssODN templates in HELA cells. Black dots represent the outcome of gene editing with unprotected ssODN templates and light blue dots represents the gene editing outcomes with 5′& ′ PT ssODN templates. Each dot represents an independent experiment. All p-values were calculated using un-paired t test, * p<0.05, **p<0.01, ***p<0.001.

## References

1. Sternberg, S. H., Redding, S., Jinek, M., Greene, E. C. & Doudna, J. A. DNA interrogation by the CRISPR RNA-guided endonuclease Cas9. Nature 507, 62–67 (2014).

2. Eyquem, J. et al. Targeting a CAR to the TRAC locus with CRISPR/Cas9 enhances tumour rejection. Nature 543, 113–117 (2017).

3. DeWitt, M. A. et al. Selection-free genome editing of the sickle mutation in human adult hematopoietic stem/progenitor cells. Sci. Transl. Med. 8, 360ra134 (2016).

4. Lin, S., Staahl, B. T., Alla, R. K. & Doudna, J. A. Enhanced homology-directed human genome engineering by controlled timing of CRISPR/Cas9 delivery. eLife 3, e04766 (2014).

5. Miyaoka, Y. et al. Systematic quantification of HDR and NHEJ reveals effects of locus, nuclease, and cell type on genome-editing. Sci. Rep. 6, 23549 (2016).

6. Hustedt, N. & Durocher, D. The control of DNA repair by the cell cycle. Nat. Cell Biol. 19, 1–9 (2016).

7. Branzei, D. & Foiani, M. Regulation of DNA repair throughout the cell cycle. Nat. Rev. Mol. Cell Biol. 9, 297–308 (2008).

8. Petersen, L. N., Orren, D. K. & Bohr, V. A. Gene-specific and strand-specific DNA repair in the G1 and G2 phases of the cell cycle. Mol. Cell. Biol. 15, 3731–3737 (1995).

9. Richardson, C. D. et al. CRISPR-Cas9 genome editing in human cells occurs via the Fanconi anemia pathway. Nat. Genet. 50, 1132–1139 (2018).

10. Chen, P. J. et al. Enhanced prime editing systems by manipulating cellular determinants of editing outcomes. Cell 184, 5635-5652.e29 (2021).

11. Ferreira da Silva, J. et al. Prime editing efficiency and fidelity are enhanced in the absence of mismatch repair. Nat. Commun. 13, 760 (2022).

12. Wang, A. T. & Smogorzewska, A. SnapShot: Fanconi anemia and associated proteins. Cell 160, 354-354.e1 (2015).

13. Nakanishi, K. et al. Human Fanconi anemia monoubiquitination pathway promotes homologous DNA repair. Proc Natl Acad Sci USA 102, 1110–1115 (2005).

14. Auerbach, A. D. Fanconi anemia and its diagnosis. Mutat. Res. 668, 4–10 (2009).

15. van de Vrugt, H. J. et al. Effective CRISPR/Cas9-mediated correction of a Fanconi anemia defect by error-prone end joining or templated repair. Sci. Rep. 9, 768 (2019).

16. Kawashima, N. et al. Correction of fanconi anemia mutation using the crispr/cas9 system. Blood 126, 3622–3622 (2015).

17. Román-Rodríguez, F. J. et al. NHEJ-Mediated Repair of CRISPR-Cas9-Induced DNA Breaks Efficiently Corrects Mutations in HSPCs from Patients with Fanconi Anemia. Cell Stem Cell 25, 607-621.e7 (2019).

18. Richardson, C. D., Ray, G. J., DeWitt, M. A., Curie, G. L. & Corn, J. E. Enhancing homology-directed genome editing by catalytically active and inactive CRISPR-Cas9 using asymmetric donor DNA. Nat. Biotechnol. 34, 339–344 (2016).

19. Rio, P. et al. Targeted gene therapy and cell reprogramming in Fanconi anemia. EMBO Mol. Med. 6, 835–848 (2014).

20. Diez, B. et al. Therapeutic gene editing in CD34+ hematopoietic progenitors from Fanconi anemia patients. EMBO Mol. Med. 9, 1574–1588 (2017).

21. Wienert, B. et al. Timed inhibition of CDC7 increases CRISPR-Cas9 mediated templated repair. Nat. Commun. 11, 2109 (2020).

22. Horlbeck, M. A. et al. Compact and highly active next-generation libraries for CRISPR-mediated gene repression and activation. eLife 5, (2016).

23. Li, W. et al. MAGeCK enables robust identification of essential genes from genome-scale CRISPR/Cas9 knockout screens. Genome Biol. 15, 554 (2014).

24. Colic, M. et al. Identifying chemogenetic interactions from CRISPR screens with drugZ. Genome Med. 11, 52 (2019).

25. Yang, Y.-G., Lindahl, T. & Barnes, D. E. Trex1 exonuclease degrades ssDNA to prevent chronic checkpoint activation and autoimmune disease. Cell 131, 873–886 (2007).

26. Stetson, D. B., Ko, J. S., Heidmann, T. & Medzhitov, R. Trex1 prevents cell-intrinsic initiation of autoimmunity. Cell 134, 587–598 (2008).

27. Ablasser, A. et al. TREX1 deficiency triggers cell-autonomous immunity in a cGAS-dependent manner. J. Immunol. 192, 5993–5997 (2014).

28. Mazur, D. J. & Perrino, F. W. Identification and expression of the TREX1 and TREX2 cDNA sequences encoding mammalian 3’-->5’ exonucleases. J. Biol. Chem. 274, 19655–19660 (1999).

29. Crow, Y. J. et al. Mutations in the gene encoding the 3’-5’ DNA exonuclease TREX1 cause Aicardi-Goutières syndrome at the AGS1 locus. Nat. Genet. 38, 917–920 (2006).

30. Fye, J. M., Orebaugh, C. D., Coffin, S. R., Hollis, T. & Perrino, F. W. Dominant mutation of the TREX1 exonuclease gene in lupus and Aicardi-Goutieres syndrome. J. Biol. Chem. 286, 32373–32382 (2011).

31. Rice, G. et al. Heterozygous mutations in TREX1 cause familial chilblain lupus and dominant Aicardi-Goutieres syndrome. Am. J. Hum. Genet. 80, 811–815 (2007).

32. Mohr, L. et al. ER-directed TREX1 limits cGAS activation at micronuclei. Mol. Cell 81, 724-738.e9 (2021).

33. Bochkareva, E., Frappier, L., Edwards, A. M. & Bochkarev, A. The RPA32 subunit of human replication protein A contains a single-stranded DNA-binding domain. J. Biol. Chem. 273, 3932–3936 (1998).

34. Terrazas, M., Alagia, A., Faustino, I., Orozco, M. & Eritja, R. Functionalization of the 3’-ends of DNA and RNA strands with N-ethyl-N-coupled nucleosides: a promising approach to avoid 3’-exonuclease-catalyzed hydrolysis of therapeutic oligonucleotides. Chembiochem 14, 510–520 (2013).

35. Möller, L. et al. Recursive Editing improves homology-directed repair through retargeting of undesired outcomes. Nat. Commun. 13, 4550 (2022).

36. Canny, M. D. et al. Inhibition of 53BP1 favors homology-dependent DNA repair and increases CRISPR-Cas9 genome-editing efficiency. Nat. Biotechnol. 36, 95–102 (2018).

37. Orebaugh, C. D. et al. The TREX1 C-terminal region controls cellular localization through ubiquitination. J. Biol. Chem. 288, 28881–28892 (2013).

38. Miyazaki, T. et al. The 3’-5’ DNA exonuclease TREX1 directly interacts with poly(ADP-ribose) polymerase-1 (PARP1) during the DNA damage response. J. Biol. Chem. 289, 32548–32558 (2014).

39. Renaud, J.-B. et al. Improved Genome Editing Efficiency and Flexibility Using Modified Oligonucleotides with TALEN and CRISPR-Cas9 Nucleases. Cell Rep. 14, 2263–2272 (2016).

40. Liang, X., Potter, J., Kumar, S., Ravinder, N. & Chesnut, J. D. Enhanced CRISPR/Cas9-mediated precise genome editing by improved design and delivery of gRNA, Cas9 nuclease, and donor DNA. J. Biotechnol. 241, 136– 146 (2017).

41. Shy, B. R. et al. High-yield genome engineering in primary cells using a hybrid ssDNA repair template and small-molecule cocktails. Nat. Biotechnol. (2022) doi:10.1038/s41587-022-01418-8.

42. Iyer, S. et al. Efficient Homology-Directed Repair with Circular Single-Stranded DNA Donors. The CRISPR Journal (2022) doi:10.1089/crispr.2022.0058.

43. Rios, X. et al. Stable gene targeting in human cells using single-strand oligonucleotides with modified bases. PLoS ONE 7, e36697 (2012).

44. Wienert, B., Shin, J., Zelin, E., Pestal, K. & Corn, J. E. In vitro-transcribed guide RNAs trigger an innate immune response via the RIG-I pathway. PLoS Biol. 16, e2005840 (2018).

45. Clement, K. et al. CRISPResso2 provides accurate and rapid genome editing sequence analysis. Nat. Biotechnol. 37, 224–226 (2019).

