## Supplementary_figures for "TREX1 restricts CRISPR-Cas9 genome editing in human cells"

#### Supplementary figure 1

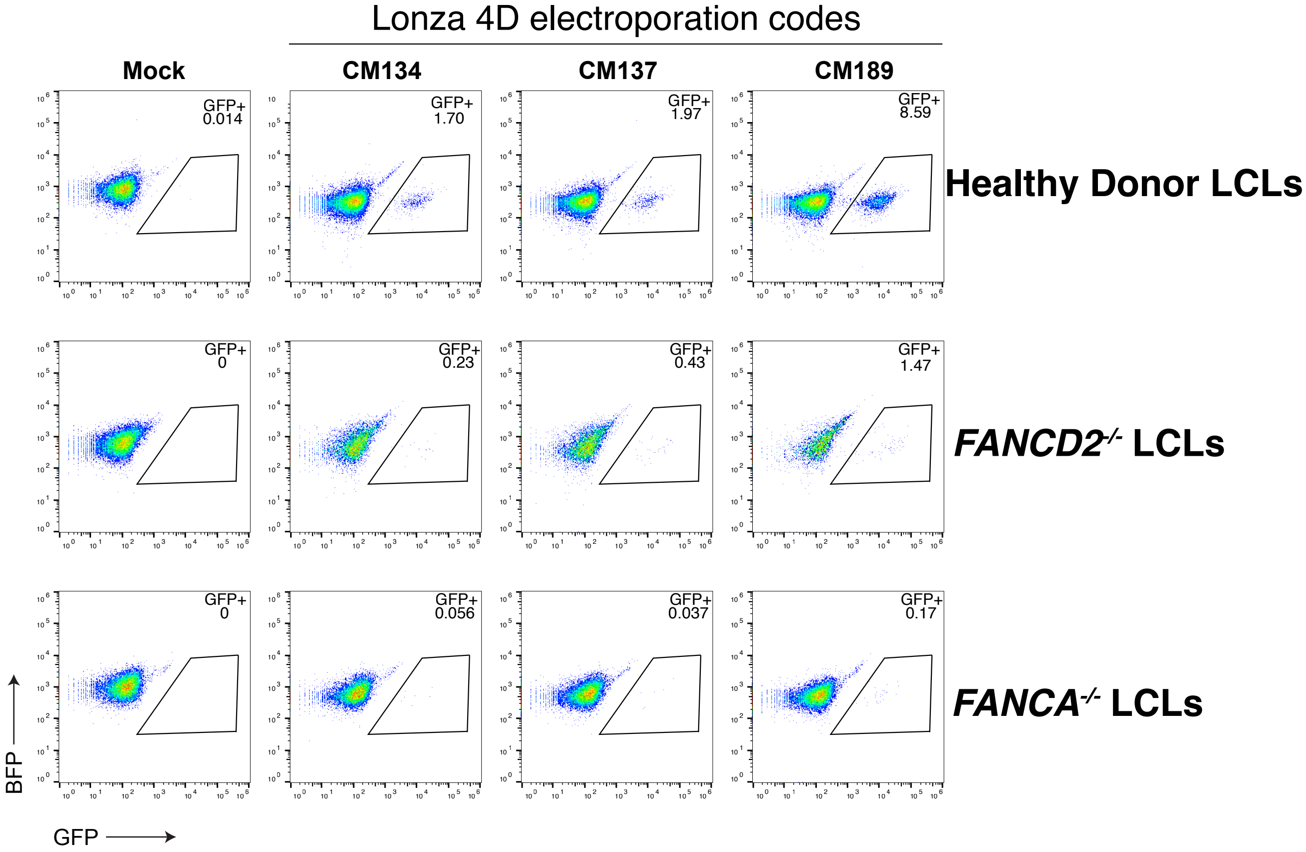

### Supplementary figure 2

BD FACSDiva 9.0.1

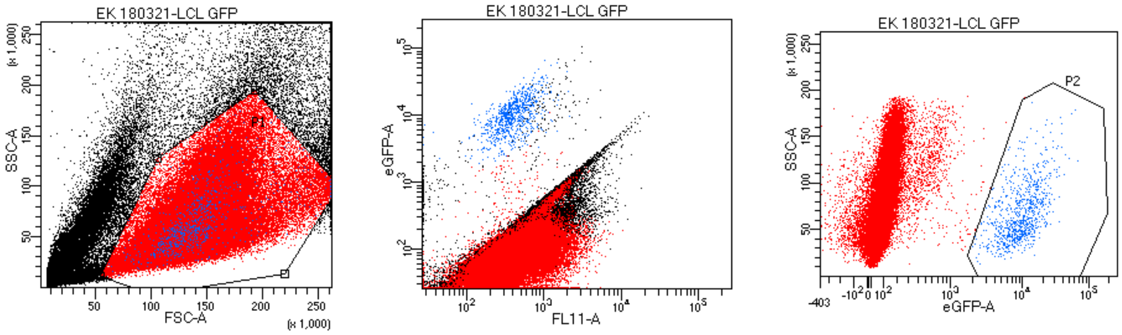


Tube: LCL GFP

| Population | #Events | %Parent | %Total |
|---|---|---|---|
| All Events | 200,000 | #### | 100.0 |
| P1 | 148,888 | 74.4 | 74.4 |
| P2 | 814 | 0.5 | 0.4 |

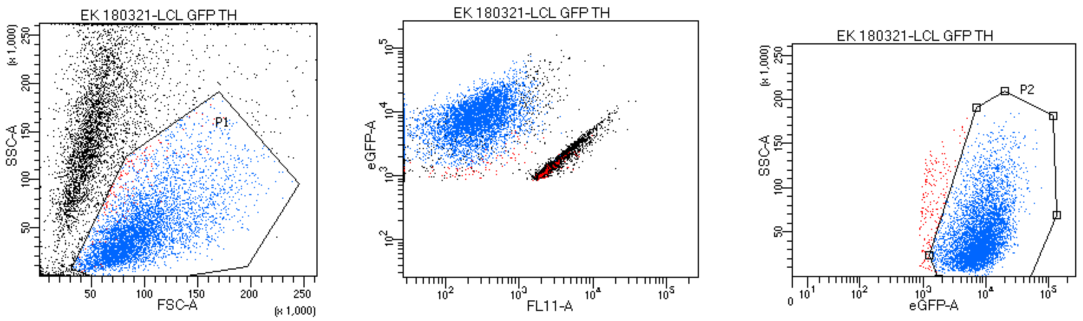


Tube: LCL GFP 2nd run

| Population | #Events | %Parent | %Total |
|---|---|---|---|
| All Events | 496 | #### | 100.0 |
| P1 | 269 | 54.2 | 54.2 |
| P2 | 215 | 79.9 | 43.3 |

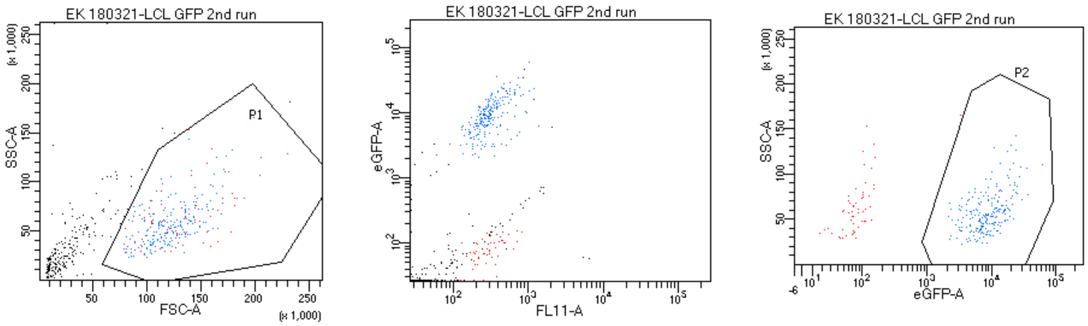


Tube: LCL GFP TH

| Population | #Events | %Parent | %Total |
|---|---|---|---|
| All Events | 10,000 | #### | 100.0 |
| P1 | 5,416 | 54.2 | 54.2 |
| P2 | 5,230 | 96.6 | 52.3 |

**Supplementary figure 3**

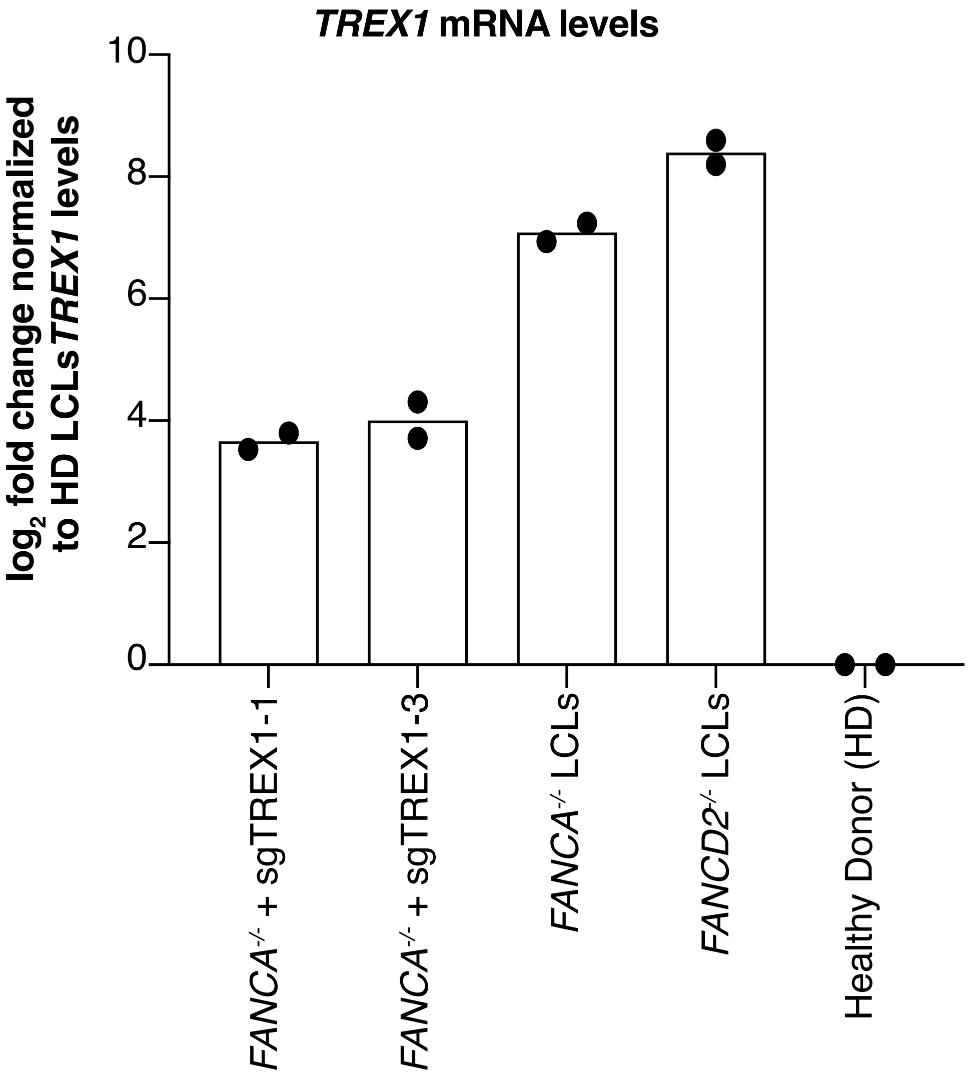

**Supplementary figure 4**

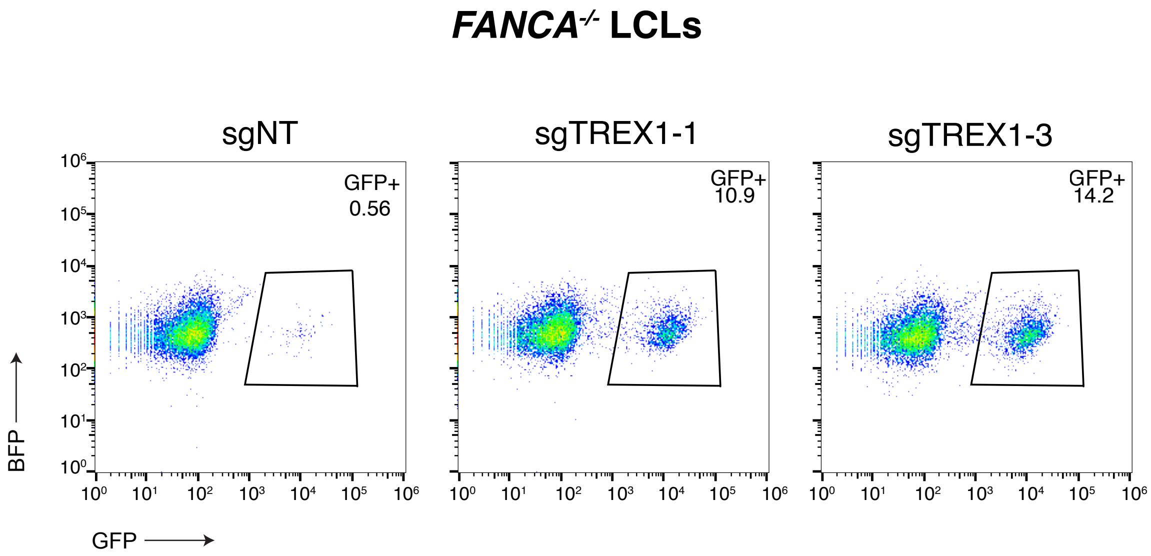

### Supplementary figure 5

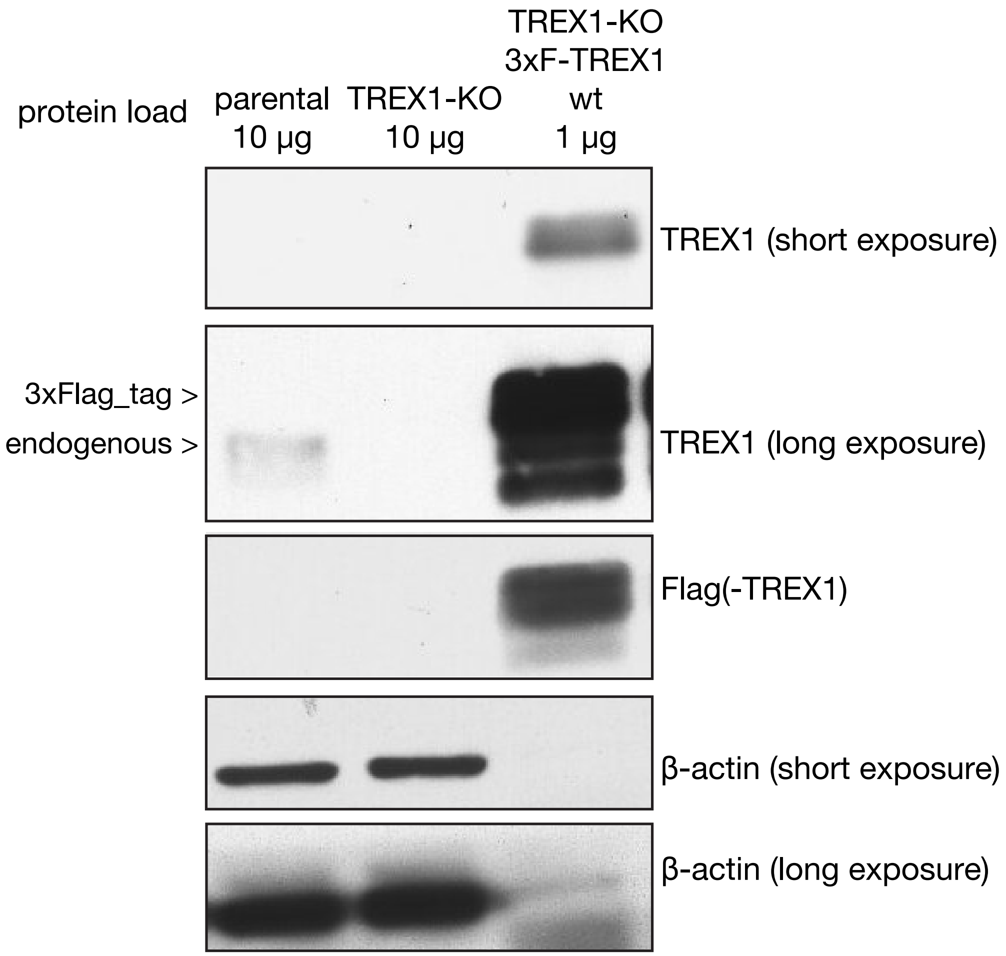

**Supplementary figure 6**

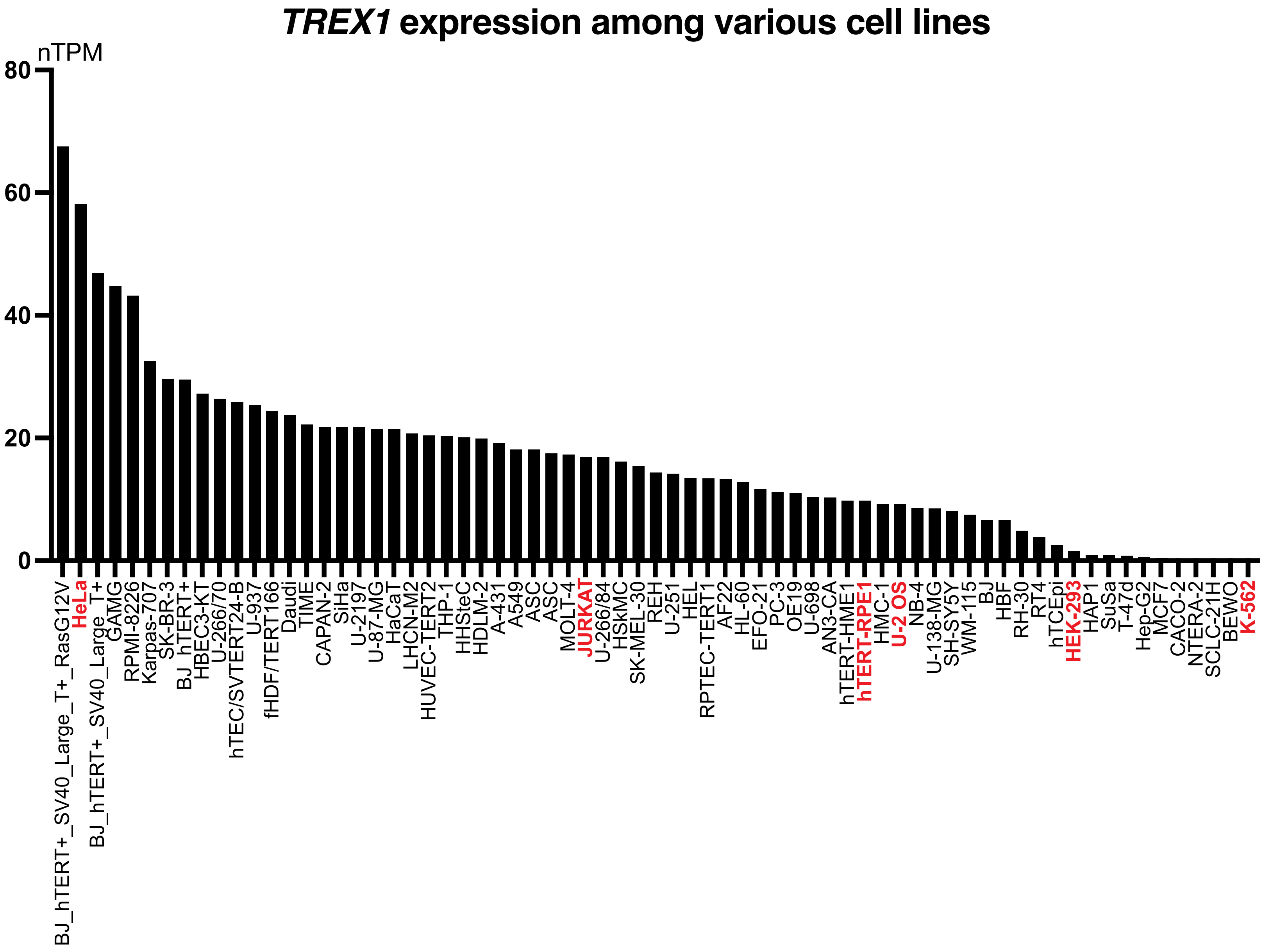

**Supplementary figure 7**

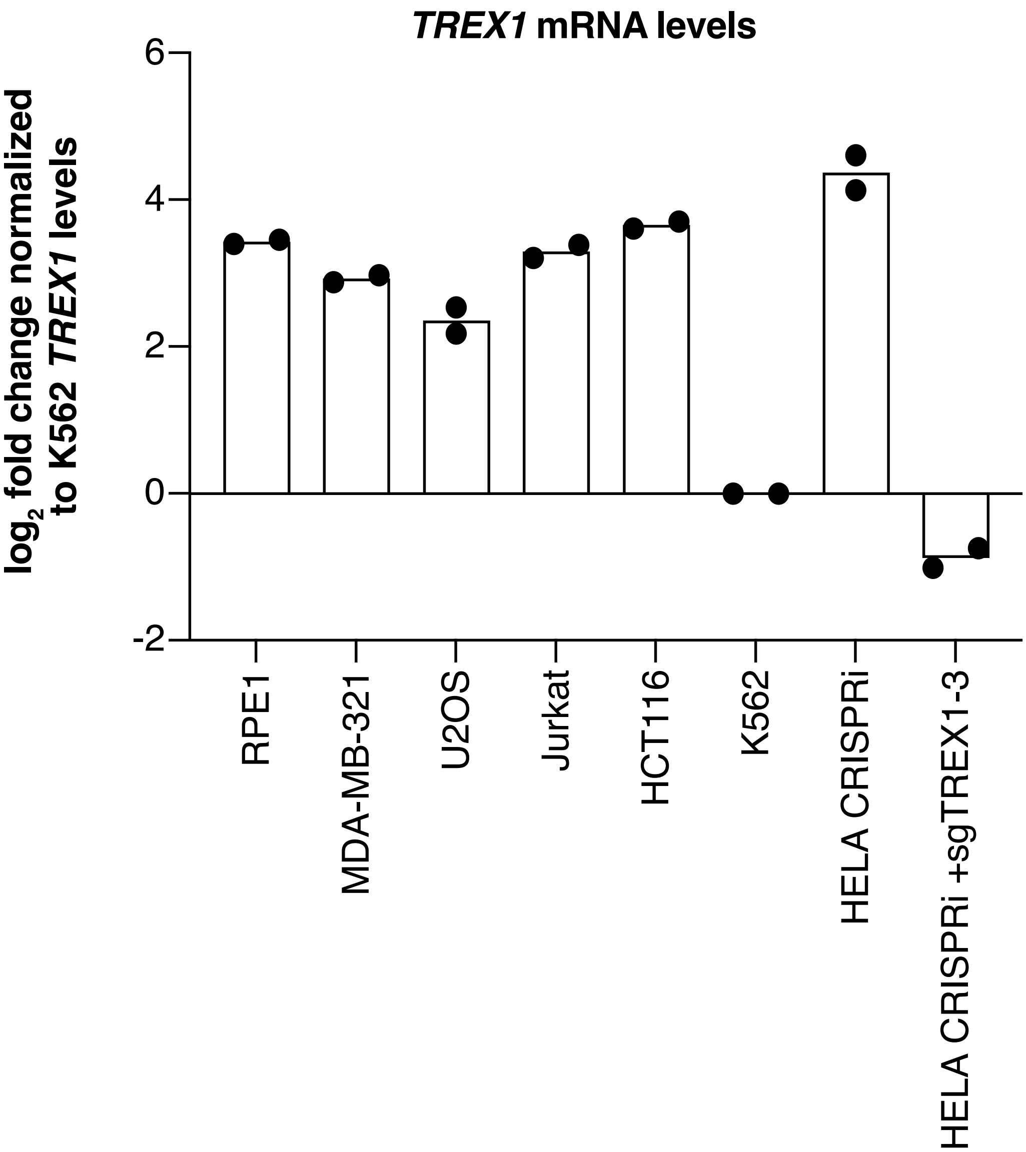

**Supplementary figure 8**

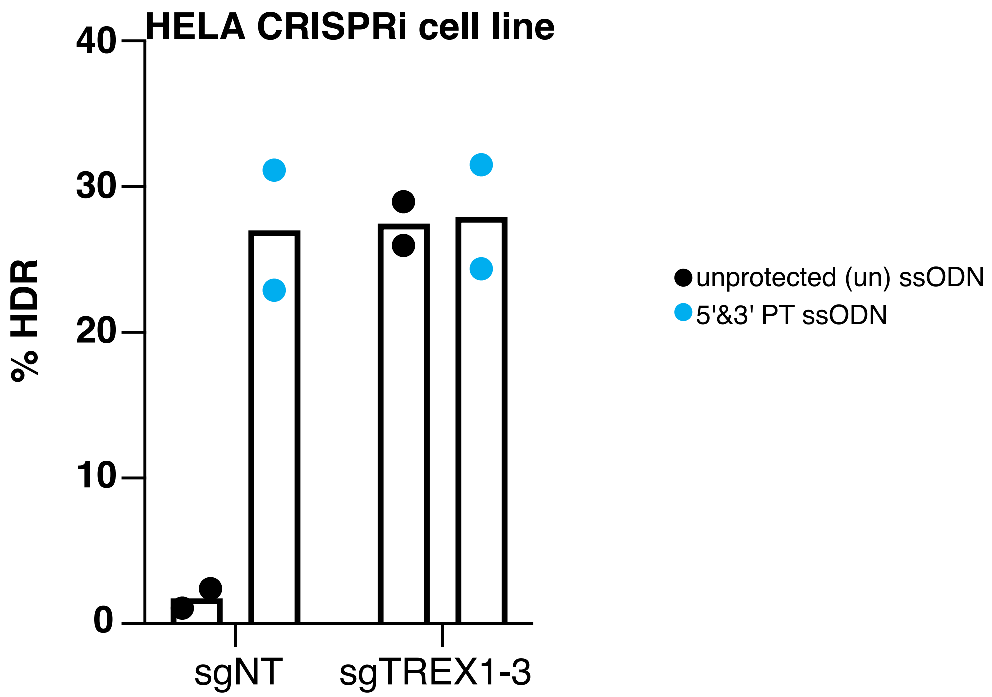

**Supplementary figure 9**

HBB

TGAGGAGAAGTCTGCCGTTACTGCCCTGTGGGCAAGG **(wild type)**
TG**TA**GAGAAGTCTGC**G**GTTACTGCCCTGTGGGCAAGG **(intended edit)**

CXCR4

GACCTCCTCTTTGTCATCACGCTTC **(wild type)**
GA**CATGT**TCTTTGTCATCACGCTTC **(intended edit)**

FANCD2

TGACCTACTGATAGAGAATACTTCAC **(wild type)**
TGA**T**CTACTGATAGAGAATAC–TCAC **(intended edit)**

UROS

GGAAGCAGCAGAGTTATGTTTGGAG **(wild type)**
GGAAGCAGCAGAGTTAT**TAG**GTTTGGAG **(intended edit)**

ABCA3

AACCTGCTTCAGAGACTCAGGGGCATCC **(wild type)**
AACCTGCTTCAGAGACTC**––**GGGCATCC **(intended edit)**

**Supplementary figure 10**

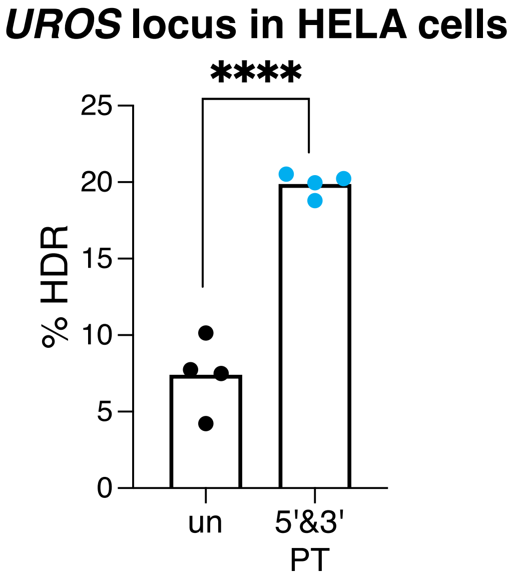
